# Peptidisc-assisted hydrophobic clustering towards the production of multimeric and multispecific nanobody proteins

**DOI:** 10.1101/2024.12.31.630897

**Authors:** Yilun Chen, Franck Duong van Hoa

## Abstract

Protein multimerization is a powerful engineering strategy for enhancing structural stability, diversity and functional performance. Typical methods to cluster proteins include tandem linking, fusion to self-assembly domains and cross-linking. We present here an approach that leverages the peptidisc membrane mimetic to stabilize hydrophobic-driven protein associations. We apply the method to nanobodies (Nbs), effective substitutes to antibodies due to their production efficiency, cost effectiveness, and lower immunogenicity, and we demonstrate the formation of multimeric assemblies termed “polybodies” (Pbs). Starting with Nbs directed against the green fluorescent protein (GFP), we produce Pbs that display increased affinity for GFP due to the avidity effect. The benefit of avidity in affinity-based assays is also demonstrated using moderate-affinity Nbs against human serum albumin. With the same auto-assembly principle, we produce bispecific and auto-fluorescent Pbs, validating our method as a versatile and general engineering strategy to generate multispecific and multifunctional protein entities. Peptidisc-assisted hydrophobic clustering thus expands the protein engineering toolbox to broaden the scope of protein multimerization in life sciences.

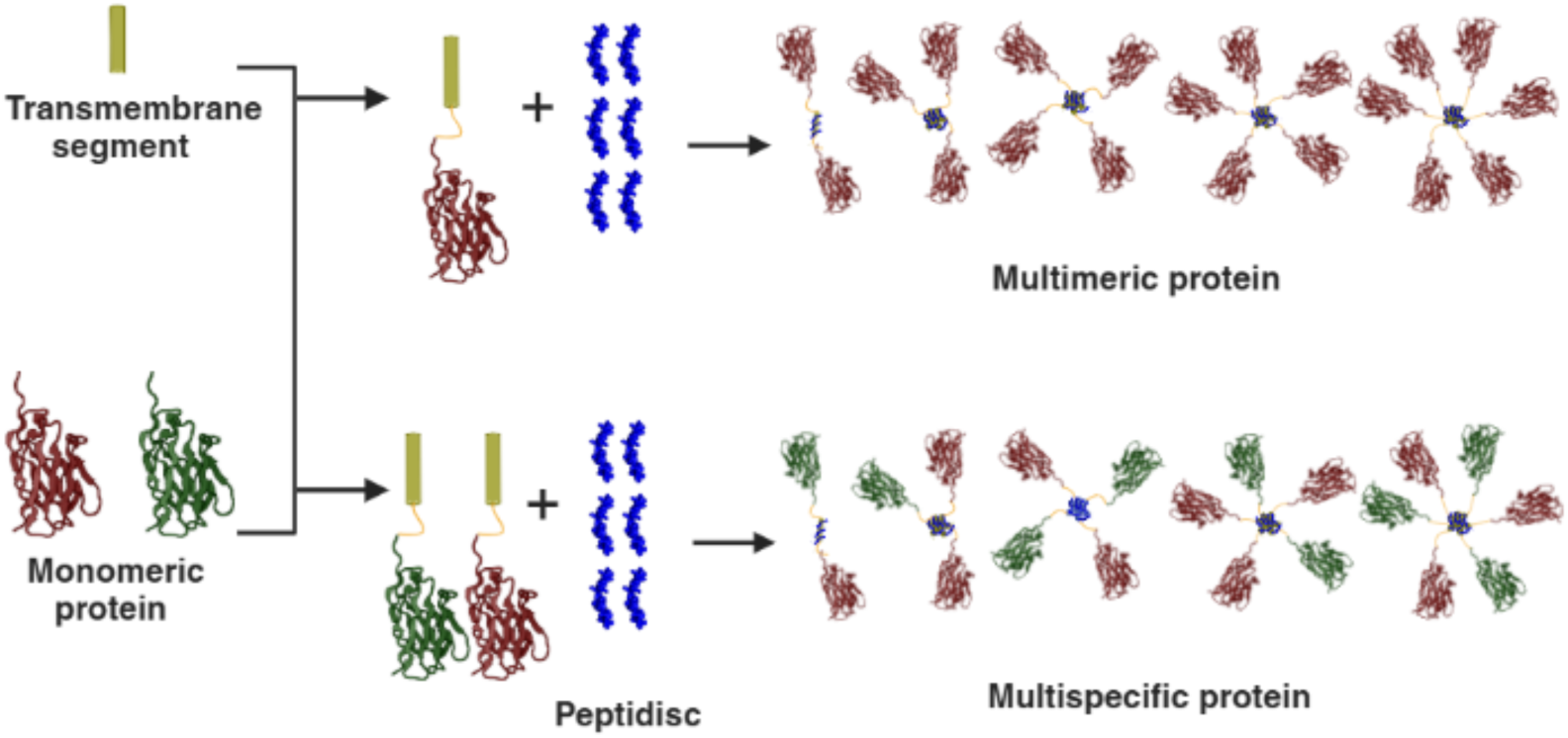

## Introduction

Approximately 30-35% of cellular proteins in nature are oligomeric, comprising either identical subunits (homo-oligomeric) or different subunits (hetero-oligomeric) ^1,2^. This prevalence is advantageous since for example multimerization allows proteins to form larger quaternary structures without increasing genome size ^3^. Additionally, the reduced surface area of each monomer within the multimeric complex enhances stability by providing protection against degradation and denaturation ^3^. Other benefits of oligomerization include gain of function, allosteric regulation of enzyme activity, and cooperative binding properties ^4^. Consequently, methods are developed to artificially multimerize proteins to achieve stability and biological function otherwise untenable with the individual monomeric units.

Current strategies for protein multimerization fall into three classes: tandem linking, self-assembly, and crosslinking, with the first two being the most commonly employed ^5^. In the tandem linking method, multiple copies of identical or distinct protein sequences are connected in a linear arrangement by joining their N– and C-termini with a linker of choice ^5,6^. These linkers are typically classified as flexible, rigid, or cleavable, depending on the specific application ^6^. In the self-assembly process, proteins are fused to self-associating peptides or heterologous scaffold proteins that have a natural tendency to oligomerize, such as the C4-binding protein oligomerization domain (C4bp, heptamer) or the ferritin (24-mer) ^7–10^. The forces driving self-assembly commonly include Van der Waals and electrostatic interactions, hydrogen bonds, inter-chain disulfide bonds, and hydrophobic effect ^5,10,11^.

Membrane proteins, especially those containing α-helical transmembrane segments (TMS), are typically kept in solution using detergents at concentrations above their critical micelle concentration (CMC) ^12^. These hydrophobic proteins have a natural tendency to self-associate in solution, especially when the detergent concentrations approach the CMC. Below the CMC, membrane proteins lose solubility and irreversibly aggregate. Based on this observation, we thought of a protein multimerization strategy that leverages the clustering properties of membrane proteins while maintaining their water solubility after removal of detergent. The method we present involves fusing a protein of interest to a TMS, creating a thermodynamically favorable hydrophobic force driving self-association. The amphipathic peptidisc is then utilized to stabilize these oligomeric assemblies while maintaining their water-solubility ^13^.

In this report, we use nanobodies (abbreviated Nbs or sometimes nAbs) as a model to validate our peptidisc-assisted protein clustering method. Originally discovered in camels, Nbs are derived from heavy-chain-only antibodies and offer several advantages over conventional antibodies. These advantages include structural simplicity, strict monomeric behavior, low immunogenicity, ease of production, high stability, and the ability to access hidden epitopes ^14–18^. Most importantly, their strictly monomeric nature, combined with the relatively small size (∼15 kDa), makes them ideal candidates for multimerization strategies aimed at boosting functional affinity through the avidity effect, as well as for changing their overall functionality or specificity. To date, Nbs have been multimerized using both flexible and rigid linkers to form homo– and hetero-oligomers, with Caplacizumab—the first FDA-approved Nb drug—being a prominent example ^19–25^. Other approaches involve self-assembly, leveraging self-associating peptides (e.g., cartilage oligomeric matrix protein, C4bp, mutated p53 tetramerization domain, tetrabrachion, and vasodilator-stimulated phosphoprotein) or self-polymerizing proteins (e.g., Fc regions, streptavidin, verotoxin 1 B-subunit, ferritin, and lumazine synthase) ^7,26–34^. Chemical and enzymatic methods, such as Sortase A-mediated ligation and Sulfo-SMCC crosslinking, have also been explored ^35,36^. A recent review summarizes the principles, advantages, and limitations of these strategies ^37^. All these methods typically rely on hydrophilic proteins or domains to promote multimerization, and none capitalize on the hydrophobic effect.

In this study, Nbs are fused to TMS and stabilized with the peptidisc scaffold, hereafter referred to as “polybodies” (Pbs). We demonstrate the successful production of anti-GFP Pbs, which exhibits avidity with an almost negligible rate of dissociation from its target. Additionally, we reformat a moderate-affinity human serum albumin (HSA) Nb into a Pb, which significantly enhances antigen detection in enzyme-linked immunosorbent assays (ELISA). Extending this concept further, we report the capture of multiple GFP and HSA nanobodies in the same polybody (GFP/HSA Pbs), as well as the incorporation of the GFP along the HSA nanobody to create auto-fluorescent Pbs. Our multimerization method thus emerges as a versatile strategy for generating multispecific and multifunctional protein assemblies.

## Material and Methods

### Plasmids and reagents

The *E. coli* strain BL21 (DE3) and the plasmid encoding his-tagged GFP (Trc99a-GFPhis_6_) are sourced from our laboratory collection. The pOPINE GFP nanobody was a gift from Brett Collins (Addgene plasmid #49172; http://n2t.net/addgene:49172; RRID:Addgene_49172). The pET26b_Nb.b201 HSA nanobody was a gift from Andrew Kruse (Addgene plasmid #131404; http://n2t.net/addgene:131404; RRID:Addgene_131404). Plasmids encoding His-tagged GFP nanobody-TMS (pBAD22-his_6_GFPNbTMS), His-tagged HSA nanobody-TMS (pBAD22-his_6_HSANbTMS) and His-tagged GFP-TMS (pBAD22-his_6_GFPTMS) were constructed using the PIPE method by joining the corresponding protein sequences to the linker (N_ter_-NPFPPMMDTLQNM-C_ter_) and the transmembrane segment (TMS) (N_ter_-ATRPALWILLVAIILMLVWLVR-C_ter_) ^38^. All constructs were verified by DNA sequencing (Genewiz). Peptidisc (NSPr and Biotin-NSPr, purity >90%) was obtained from Peptidisc Biotech. Detergent N-dodecyl-β-d-maltoside (DDM) was purchased from Anatrace. Nickel-chelating agarose was sourced from Qiagen. Superdex 200 resin was from GE Healthcare. Tryptone, yeast extract, NaCl, imidazole, Tris-base, acrylamide 40%, bis-acrylamide 2%, TEMED, and bovine serum albumin (BSA) were acquired from Bioshop Canada. Isopropyl β-d-1-thiogalactopyranoside (IPTG) was purchased from Bio Basic. Arabinose and ampicillin were obtained from GoldBio. Human serum albumin (HSA) was procured from MilliporeSigma. Kanamycin and all other chemicals were obtained from Fisher Scientific.

### Membrane protein and soluble protein expression and purification

The expression of His-tagged GFP nanobody-TMS and His-tagged HSA nanobody-TMS was conducted in *E. coli* BL21(DE3) in LB medium supplemented with 100 μg/mL ampicillin. The cell culture was propagated to the exponential growth phase (OD_600nm_ ∼ 0.6) at 37°C, and protein synthesis was induced by the addition of 0.2% of arabinose. Further growth was carried out at 20°C for 20 hours. Cells were harvested by low-speed centrifugation (6,000*g*, 8 min) and resuspended in buffer TSB (50 mM Tris-HCl pH 7.8, 100 mM NaCl, 100 mM Na_2_CO₃) supplemented with 1 mM phenylmethylsulfonyl fluoride (PMSF). All subsequent steps were carried out at 4°C. Cells were lysed through a microfluidizer (Microfluidics; 3 passes at 15,000 psi). Unbroken cells were removed by low-speed centrifugation (6,000*g*, 10 min). The crude membrane fraction was isolated by ultracentrifugation (100,000*g*, 45 min, Beckman Coulter rotor Ti45). Membrane pellets were resuspended in buffer TSG (50 mM Tris-HCl pH 7.8, 100 mM NaCl, 10% glycerol) at 10 mg/mL and stored at –20°C for later use. His-tagged GFP-TMS was produced in *E. coli* BL21(DE3) at 37°C in LB medium supplemented with 100 μg/mL ampicillin. 0.2% arabinose was added during the exponential growth phase (OD_600nm_ ∼ 0.6). After 3 hours, cells were harvested and lysed as described above. His-tagged GFP nanobody and His-tagged HSA nanobody were produced as previously described ^39,40^. Expression of His-tagged GFP was conducted in *E. coli* BL21(DE3) in LB medium supplemented with 100 μg/mL ampicillin. The cell culture was propagated to the exponential growth phase (OD_600nm_ ∼ 0.6) at 37°C, and protein synthesis was induced by the addition of 1 mM IPTG for 3 hours. After 3 hours, cells were harvested and lysed as described above. The cleared cell lysate was isolated by ultracentrifugation (100,000*g*, 45 min, Beckman Coulter rotor Ti45) and stored at –20°C for later use. GFP, GFP Nbs and HSA Nbs were purified by Ni^2+^-chelating chromatography (Ni-NTA). The cleared cell lysate was passed through a hand-packed Ni-NTA gravity column (1 mL resin), and the flowthrough was discarded. Subsequently, the resin was washed with 50 mL of Ni-NTA wash buffer (TSG buffer 20 mM imidazole). Bound proteins were eluted with Ni-NTA elution buffer (TSG buffer, with 600 mM imidazole). The purity of the proteins was analyzed by SDS-PAGE. Purified proteins were stored at –20°C for further applications and analysis.

### Reconstitution of membrane proteins in peptidisc via PeptiQuick method

All steps were carried out at 4°C. For GFP nanobody-TMS and HSA nanobody-TMS, about 20 mg of crude membranes were solubilized with 1% DDM for 45 min. After the removal of insoluble aggregates by ultracentrifugation (180,000g, 15 min), the detergent extract was reconstituted into peptidisc as previously described with minor modifications ^41^. Briefly, the detergent extract was incubated with pre-equilibrated Ni-NTA resin (150 µL) for 1 hour on a tabletop rocker. The resin was then sedimented and washed with 30 column volumes (CV; 1.5 mL) of Buffer TSGD (TSG buffer, 20 mM imidazole, 0.02% DDM). The resin was then resuspended in 1 mL of buffer TSG supplemented with 1 mg of peptidisc. The resin was washed with 30 CV buffer TSG to remove excess peptidisc. The peptidisc-reconstituted proteins were eluted in Ni-NTA elution buffer (200 µL). For GFP-TMS, approximately 5 mg of crude membranes were solubilized with 1% DDM for 45 min, followed by all subsequent steps as described above. The purity and reconstitution of the proteins were analyzed by SDS-PAGE and native gel electrophoresis. Protein samples were stored at –20°C before further applications and analysis.

### Native gel electrophoresis

Equal volumes of 4% and 12% acrylamide solutions were prepared in advance, and linear gradient gels were poured manually. The cross-linking agents, TEMED and ammonium persulfate, were added immediately before gradient mixing. Plastic combs (Bio-Rad) were then inserted, and the gels were allowed to polymerize for 30 min before storage at 4°C. For clear native gel electrophoresis, the anode and cathode buffers consisted of buffer N (37 mM Tris-HCl pH 8.8, 35 mM Glycine). For blue-native gel electrophoresis, the anode buffer was replaced with buffer NA (Buffer N, 180 µM Coomassie Blue G-250), while the cathode buffer remained unchanged.

### Gel shift assay

The binding reaction consisted of a mixture (20 μL) containing 4 μg of purified GFP Pb or GFP Nb and either 4 μg or 8 μg of GFP in buffer TSG. Following a 15-minute incubation at 20°C, the assay mixtures were subjected to clear native gel electrophoresis. Before Coomassie Blue staining, the gel was scanned at the Cy2 channel (530 nm) using the Amersham Typhoon Biomolecular Imager (Cytiva).

### Size-exclusion chromatography and electron microscopy

To isolate the GFP:Pb complex, purified Pbs were mixed with a 2-fold molar excess of GFP, followed by a 15 min incubation at 20°C. The mixture was then applied to a Superdex^TM^ 200 (10/300) column pre-equilibrated with buffer TSG, with a flow rate set at 0.5 mL/min. Fractions collected from the column were analyzed on clear native gels, and those containing the complex (F1-F9) were pooled based on relative mobility (F2-F3, F4-F5, and F6-F7). Protein concentration was determined using the Bradford assay reagent (Bio-Rad), and concentrations were adjusted to 0.02 mg/ml with buffer TSG. Subsequently, the protein samples were adsorbed onto glow-discharged carbon-coated grids (Electron Microscopy Sciences) and stained with 0.75% uranyl formate (Electron Microscopy Sciences). Imaging was performed using a Talos microscope (ThermoFisher) operating at an accelerating voltage of 120 kV, with images acquired at a magnification of 45,000 and a defocus of 3 μm.

### Biolayer interferometry

Streptavidin-coated sensors (Gator Bio) were employed with the Gator® Prime instrument (Gator Bio). For the affinity measurement, GFP Pbs was reconstituted using the method described above, utilizing a peptidisc with a biotin tag. To biotinylate GFP Nb, the protein was rebuffered to buffer HSG (50 mM HEPES-NaOH, pH 7.8, 100 mM NaCl, 10% glycerol), and mixed with a 20-fold molar excess of NHS-Biotin (ApexBio). The reaction mixture was incubated at 20°C for 3 hours, followed by removal of unreacted NHS-Biotin via dialysis to Buffer TSG at 4°C for 16 hours using a 6-8 kDa MWCO dialysis bag (Spectrum). Sensors were equilibrated with 250 μL of kinetic buffer (50 mM Tris-HCl pH 7.8, 100 mM NaCl, 0.05% BSA) at 20°C for 15 min in the Max Plate (Gator Bio). Assays were conducted in the BLI 96-Flat Plate (Gator Bio). Following equilibration, sensors were exposed to 200 μL of a 5 μg/mL solution of biotinylated GFP Pbs or a 10 μg/mL solution of biotinylated GFP Nbs and subsequently washed with kinetic buffer. Interactions between GFP:PBs and GFP:Nbs were measured using 200 μL of serially diluted GFP (ranging from 100 nM to 3.125 nM). The signal from the empty reference probe was subtracted from the actual assay probes, and the collected data were analyzed using Gator® software (version 2.7.3.1013), employing the 1:1 model. For the avidity measuring, the GFP Pb was reconstituted using regular peptidisc. Biotinylation of GFP was conducted as previously described. Sensors were equilibrated with 250 μL of kinetic buffer at 20°C for 15 min in the Max Plate. Following equilibration, sensors were exposed to 200 μL of a 10 μg/mL solution of biotinylated GFP and subsequently washed with kinetic buffer. The interaction between GFP and the GFP Nb was measured using 200 μL of 20-40 nM GFP Nb solution. The interaction between GFP and Pbs was measured using 200 μL of diluted GFP Pbs (ranging from 10 μg/mL to 2 μg/mL). The signal from the empty reference probe was subtracted from the actual assay probes, and the collected data were analyzed using Gator® software.

### ELISA assays

100 μL per well of serially diluted HSA (in PBS, ranging from 4000 ng/mL to 1.9 ng/mL) was coated in a 96-well microplate (Corning) at 4°C for 18 hours. After washing twice with PBS (137 mM NaCl, 2.7 mM KCl, 10 mM Na_2_HPO_4_, 1.8 mM KH_2_PO_4_), blocking was performed using 0.5% BSA in PBS (300 μL per well) at 37°C for 1 hour. Following blocking, 100 μL of diluted HSA Pbs or HSA Nbs at 1 μg/mL were added to each well, followed by incubation at 37°C for 1 hour. The microplate was then washed five times with PBST (PBS, 0.002% tween-20), and 100 μL/well of HisProbe-HRP (ThermoFisher) at 1 μg/mL was added and incubated at 37°C for 45 min. After washing five times with PBST, 100 μL of TMB substrate solution (MilliporeSigma) was added into the wells and incubated at 20°C for 9 min. The reaction was stopped by the addition of 50 μL per well of 2 M H_2_SO4. Absorbance was measured at 450 nm using the Spark 20M Plate Reader (Tecan). The ELISA assays were performed in triplicate to establish the standard deviation. The detection limit was determined based on a cutoff value (mean of the control + 2× standard deviation) ^33^.

### GFP/HSA polybody preparation

10 mg of crude membrane containing GFP nanobody-TMS and 10 mg of crude membrane containing HSA nanobody-TMS were each solubilized separately with 1% DDM for 45 minutes. The resulting detergent extracts were then thoroughly mixed on a table rocker before proceeding with the PeptiQuick workflow, as previously described. To generate the GFP+HSA Pbs for control, GFP and HSA Pbs were individually produced from the respective membranes, following the previously outlined methods. After quantifying the protein concentration using the Bradford assay, the prepared GFP and HSA Pbs were combined to achieve a 1:1 stoichiometry ratio. The prepared Pbs mixture was verified and compared to the prepared GFP/HSA Pbs using SDS-PAGE analysis to ensure an identical appearance in-gel.

### Fluorescent HSA polybody preparation

10 mg of crude membrane containing HSA nanobody-TMS and 5 mg of crude membrane containing GFP-TMS were separately solubilized using 1% DDM for 45 minutes. The resulting detergent extracts were thoroughly mixed on a table rocker before proceeding with the PeptiQuick method. To generate the mixture containing HSA Pbs and poly-GFP as a control, poly-GFP and HSA Pbs were individually produced. After quantifying the protein concentrations with the Bradford assay, poly-GFP and HSA Pbs were combined to achieve a 2:1 stoichiometry ratio. The prepared mixture was verified and compared to the fluorescent HSA Pbs using SDS-PAGE analysis to ensure identical appearance in-gel.

### Dot-blot assay

An approximately 4 cm by 3 cm piece of nitrocellulose membrane (Bio-Rad) was cut and placed on a clean, flat surface. 2 μL of serially diluted HSA (in PBS, ranging from 5 mg/mL to 0.13 mg/mL) was dispensed onto individual circles on the membrane. For GFP/HSA bispecific Pbs verification experiments, the GFP Nb was serially diluted (in PBS, ranging from 0.25 mg/mL to 0.015 mg/mL) and immobilized on each membrane as a positive control. The membranes were allowed to air-dry for 15 minutes and then blocked for 1 hour in 10 mL of blocking solution (PBS, 5% milk) at 20°C with gentle rocking. After blocking, HSA Pbs, GFP Pbs, GFP+HSA Pbs mixture, or GFP/HSA Pbs were added to 3 mL of incubation solution (PBS, 0.5% milk) at a final concentration of 20 μg/mL and incubated with the membrane for 1 hour at 20°C with gentle rocking. Following five washes with 10 mL of PBST, 5 mL of GFP at 3 μg/mL (in PBS) was added and incubated with each membrane for 15 minutes at 20°C with gentle rocking. After five additional washes with PBST, blot imaging was performed using the Amersham Typhoon Biomolecular Imager at the Cy2 channel. For fluorescent HSA Pbs verification experiments, after blocking, HSA Pbs, poly-GFP, HSA Pbs+poly-GFP mixture, or fluorescent HSA Pbs were added to 3 mL of incubation solution at a final concentration of 20 μg/mL and incubated with the membrane for 1 hour at 20°C with gentle rocking. Following five washes with 10 mL of PBST, blot imaging was performed directly using the Amersham Typhoon Biomolecular Imager at the Cy2 channel.

## Results

### Principle of protein multimerization

The strategy for protein polymerization is illustrated in **Fig. 1**, using the nanobody (Nb) as model protein. In this workflow, a his-tagged Nb of interest is attached to a sequence encoding for a hydrophobic transmembrane segment (TMS), resulting in an artificial membrane protein expressed in the *E. coli* envelope. The Nb-TMS protein is then extracted from the membrane fraction using a non-ionic detergent (N-dodecyl-β-D-maltoside; DDM) and subsequently immobilized on the Ni-NTA resin. The DDM micelle is then washed away while being replaced by the peptidisc scaffold following the established PeptiQuick protocol ^41^. Herein, the TMS enables hydrophobic associations between Nb proteins based on a thermodynamically favourable driving force. These hydrophobic associations are stabilized by the amphipathic peptidisc peptide whose hydrophilic side maintains contact with the aqueous environment, while its hydrophobic face sequesters protein interactions at TMS junctions. As a result, the polymeric Nb remains water-soluble.

**Figure 1:**
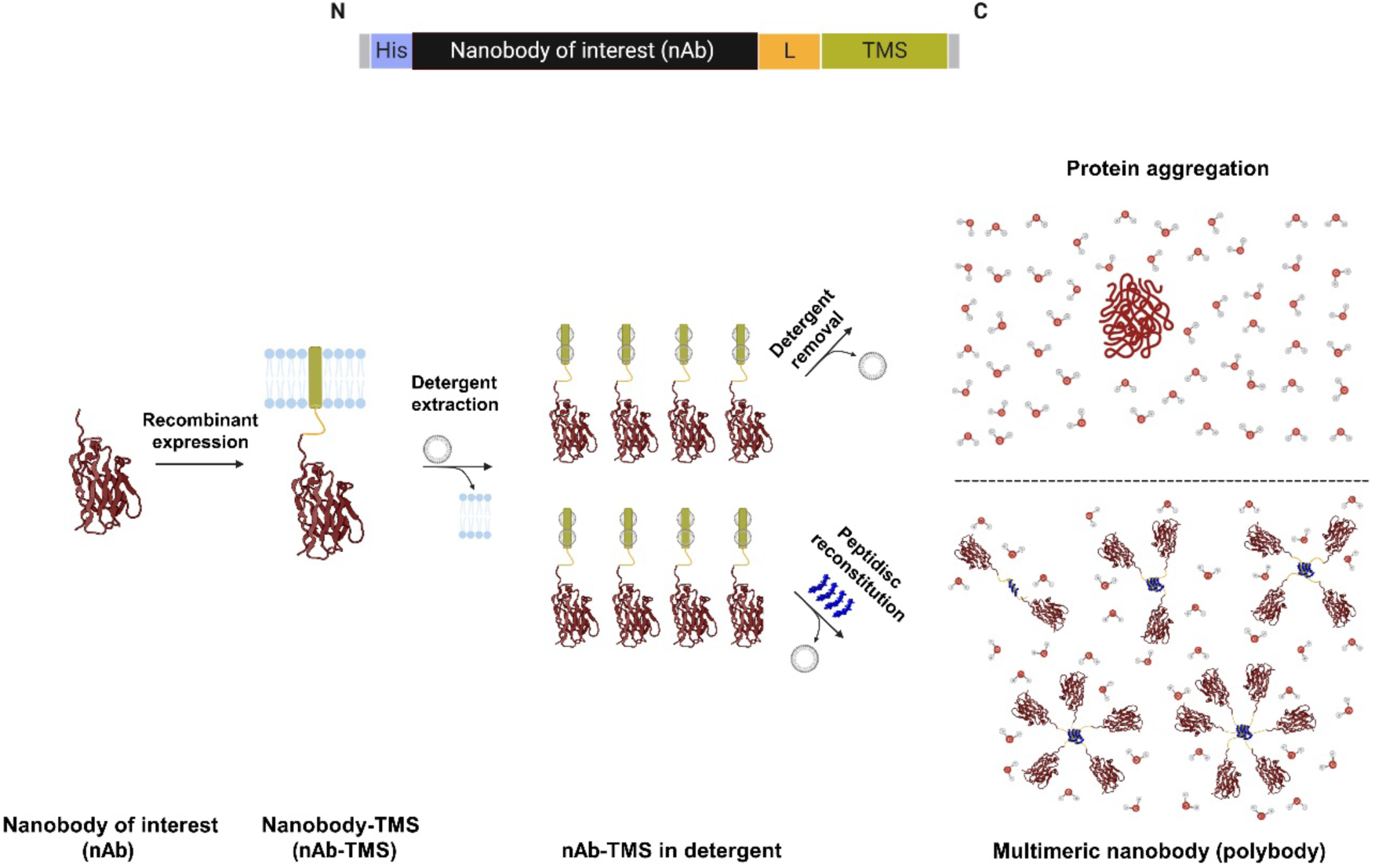
Design and principle of peptidisc-assisted protein multimerization. Demonstrated using the model system of this study: nanobodies (nAbs). Top: Primary sequence of the design. N-terminal (N), C-terminal (C), His_6_-tag (His), linker (L) region (N_ter_-NPFPPMMDTLQ NM-C_ter_), transmembrane segment (TMS) (N_ter_-ATRPALWILLVAIILMLVWLVR-C_ter_). Bottom: Schematic diagram illustrating the key steps for the capture of multiple nanobodies into a polybody. Figure created with BioRender.com

### Polymerization of a GFP Nb and interaction with GFP

To develop the polybody workflow, we employed a Nb anti-GFP ^39^. The GFP Nb sequence was fused to the TMS, resulting in a membrane protein with an estimated molecular weight of 17.7 kDa **(Fig. 2a)**. The purified protein was reconstituted in peptidisc, producing GFP Pbs with an approximate yield of 200 µg per litre of cells, which was deemed sufficient to perform initial analysis. We employed clear-native gel electrophoresis to assess the Pbs oligomeric state (**Fig. 2b**, lane 3). On this type of gel, the GFP Pbs migrated as multiple bands, corresponding to the different Nb oligomers stabilized in peptidisc. We then tested the ability of the GFP Pbs to bind and thereby alter the migration of the GFP protein using a gel shift assay (**Fig. 2b**). The gel shift due to the interaction of the Pbs with the GFP protein was visible upon Coomassie blue staining of the gel (**Fig. 2b left,** compare lane 3 to lane 5) and evident when the same gel was scanned at 530 nm to detect the fluorescent GFP (**Fig. 2b right,** compare lane 3 to lane 5). Next, GFP was mixed with Pbs and passed through a Superdex 200 gel filtration column **(Fig. 2c)**. The size-exclusion chromatography profiles showed three major peaks eluting at ∼ 10.6 mL, 16.9 mL, and 22.6 mL, respectively, with no elution observed in the void volume (∼8.0 mL), indicating the absence of large aggregates in the sample. Analysis of the fractions collected from the first peak (F1-F9) on the clear-native gel confirmed the separation of the GFP:Pbs complex from excess GFP, which appeared in the second peak (F11). The detection of the GFP:Pbs association by native gel electrophoresis and gel filtration chromatography suggests that the formation of the complex occurs efficiently. To assess the quality and stability of the Pbs another way, fractions from the gel filtration above were pooled and analyzed by negative stain electron microscopy (EM) **(Fig. 2c)**. As expected, the size of the protein particles progressively decreased with the fractionation. Importantly, no large aggregates were observed, affirming the quality and stability of the particles obtained.

**Figure 2:**
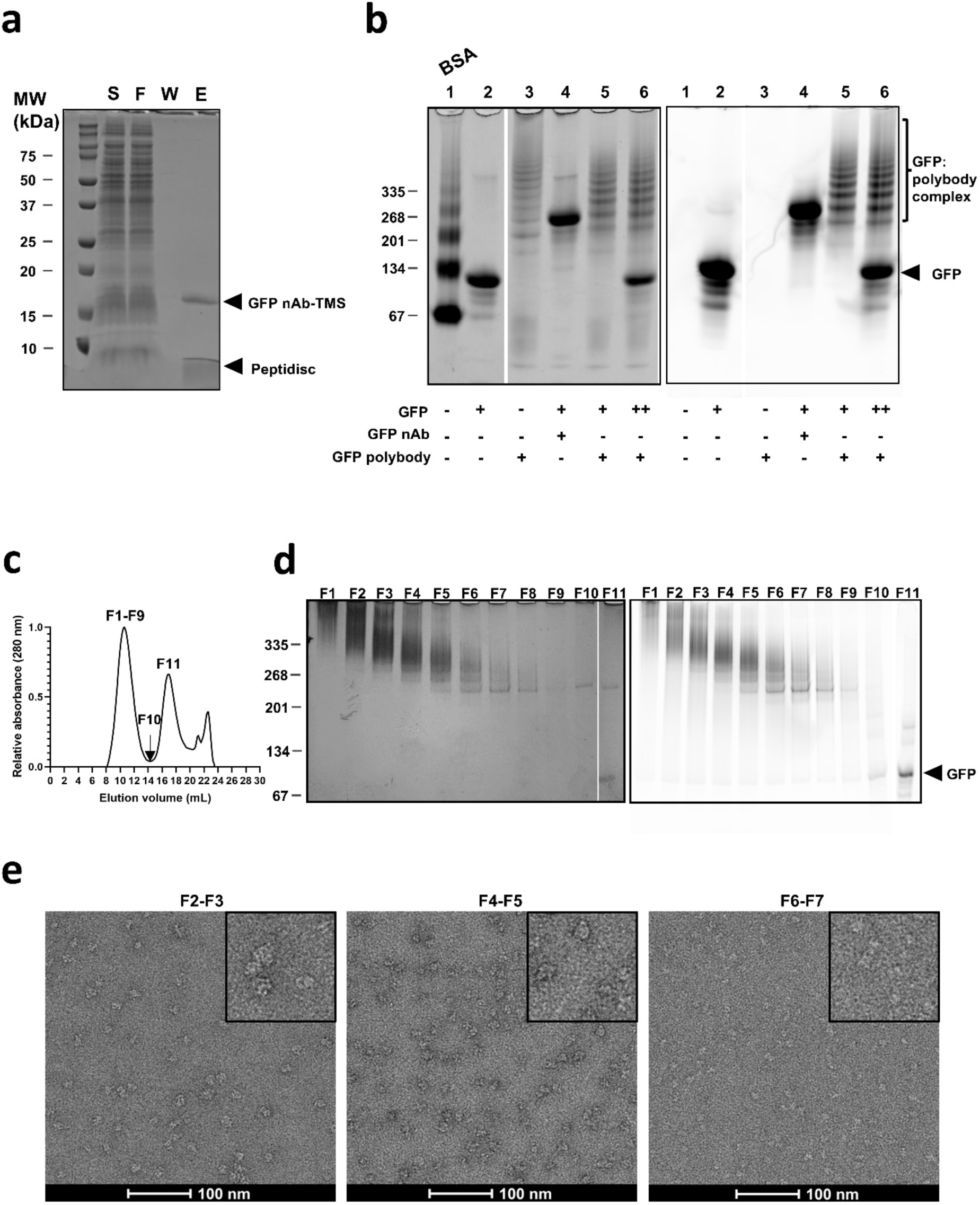
GFP polybody preparation and characterization. **(a)** SDS-PAGE analysis of the IMAC-purified and peptidisc reconstituted GFP Nb-TMS protein followed by Coomassie Blue staining of the gel. S: Starting materials (detergent extract); FT: proteins in the flow-through; W: Washing step; E: proteins eluted after peptidisc reconstitution. **(b)** Gel shift assay on a 4-12% clear native gel. 4 μg of GFP polybody (lane 3) were incubated with 4 μg and 8 μg of GFP (lane 5 & 6). The gel was first scanned at 530 nm (right), then stained with Coomassie Blue (left). The position of the GFP:polybody complex is indicated, with the GFP:GFP Nb complex shown as reference (lane 4). BSA was used as a molecular weight reference in lane 1 of the native gel. **(c)** Size-exclusion chromatography of the GFP:polybody complex on a Superdex 200 (10/300) column. Absorbance was monitored and recorded at 280 nm. **(d)** The collected SEC fractions from the first and second peaks were analyzed on 4-12% clear native gel. The gel was first scanned at 530 nm (right), then silver-stained (left). **(e)** Negative-stain EM analysis of the indicated fractions (left, fractions 2-3; middle, fractions 4-5; right, fractions 6-7).

### Binding kinetics of the GFP:polybody associations

We determined the binding kinetics of the GFP Pbs to GFP using bio-layer interferometry (BLI). In the first setup, the GFP Nbs were biotinylated with NHS-biotin and the GFP Pbs were prepared with a peptidisc containing a biotin moiety. The Nbs and Pbs preparations were immobilized on streptavidin-coated probes to monitor their interactions with the serially diluted GFP analyte **(Fig. 3a)**. This setup allowed us to monitor the binding of individual GFP molecules to each independent binding site on the GFP Pbs, enabling simple curve fitting to a 1:1 binding model for affinity determination. Both the GFP Nbs and Pbs displayed homogeneous binding patterns, as evidenced by the BLI signal increasing proportionally with GFP concentration. After subtracting signal from a reference probe (no ligand, maximum analyte concentration, **Supplementary Figure 1**), the data were fit to a 1:1 binding model. The calculated K_d_ values for the Nb and Pbs were comparable, 0.75±0.09 nM and 2.72±1.60 nM respectively **(Fig. 3b)**. The K_d_ value for the GFP Nbs aligns with previous reports ^39^, and the similar K_d_ values obtained with the Pbs indicate that the binding properties are preserved upon polymerization. The larger standard deviation obtained with the Pbs is likely attributed to the size heterogeneity due to different oligomeric states but this heterogeneity did not prevent fitting the data to the 1:1 model.

**Figure 3:**
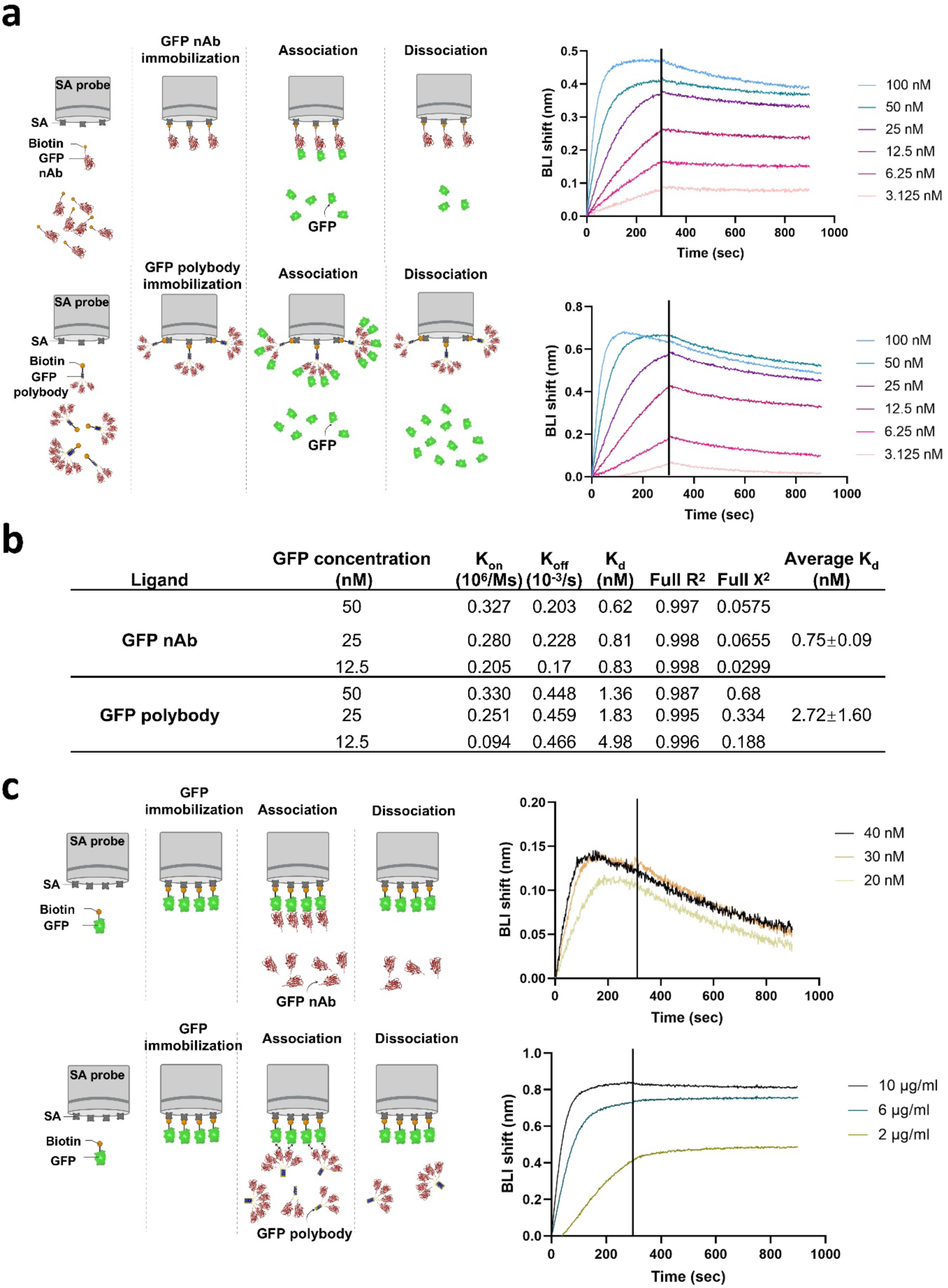
Binding kinetics of the GFP polybody using bio-layer interferometry. **(a)** Top: association and dissociation measurements between the immobilized GFP nanobody and GFP. Bottom: association and dissociation measurements between the immobilized GFP polybody and GFP. Signal from the reference probe has been subtracted. Data from all GFP dilutions (100 nM to 3.125 nM) are shown, with three concentrations (50 nM, 25 nM, and 12.5 nM) used for K_d_ calculation. The binding curve obtained with the highest GFP concentration (100 nM) could not be fitted with the polybody, which might be due to binding site saturation. **(b)** Measured association and dissociation rates (k_on_ and k_off_) and calculated dissociation constants (K_d_) for GFP Nbs and Pbs. **(c)** Avidity-biased measurement. Top: association and dissociation measurements between the immobilized GFP and GFP Nb. Bottom: association and dissociation measurements between the immobilized GFP and Pbs. Signal from the reference probe has been subtracted. Created with BioRender.com

In the second setup, we reversed the binding orientation: the biotinylated GFP was immobilized on streptavidin-coated probes, while the Nbs and Pbs were used as analytes at varying concentrations **(Fig. 3c)**. This orientation allowed the Pbs, with multiple binding sites, to interact simultaneously with multiple immobilized GFP ligands, enabling the measurement of cumulative binding affinities through the avidity effect ^42^. As anticipated, the GFP Pbs generated a much higher signal than the Nb given its larger size. Most importantly, the Pbs exhibited a near-zero dissociation rate, which is explained by the avidity effect as documented in other studies ^43,44^. In contrast, the GFP Nb produced similar Kd values in both assay orientations (0.75 ± 0.09 nM vs 3.08 ± 0.62 nM) **(Fig. 3b and Supplementary Table 1)**, which confirms that the very low dissociation rate is due to the polymeric assembly of the Pbs. However, the Pbs data could not be accurately fit to standard 1:1, 2:1 (heterogeneous ligand), or 1:2 (bivalent analyte) binding models, making precise quantification challenging, likely due to heterogeneity in its preparation.

### Increasing the HSA nanobody affinity for ELISA application

To showcase the utility of the Pbs, we employed a Nb with a moderate affinity for its target, here the human serum albumin (HSA Nb; K_d_ ∼ 430 nM) ^40^. Following cloning, purification and reconstitution in peptidisc, the HSA Nbs assembled in oligomers characteristic of Pbs formation **(Fig. 4a)**. We then assessed the performance of the HSA Pbs using an indirect ELISA assay. To facilitate comparison, the concentration of the HSA Nbs and Pbs were estimated by SDS-PAGE analysis and adjusted accordingly **(Fig. 4b)**. Serial dilutions of the HSA protein (1.9-4000 ng/mL) were then coated onto the microplate and incubated with equivalent amounts of Nbs or Pbs. The binding was monitored using the reagent HisProbe-HRP, which recognizes the His_6_ tag **(Fig. 4c)**. In this assay, the limit of detection (LOD) for the HSA Nbs was determined at 1000 ng/mL. The LOD for the HSA Pbs was 31.25 ng/mL, representing a 32-fold enhancement in the HSA detection sensitivity.

**Figure 4:**
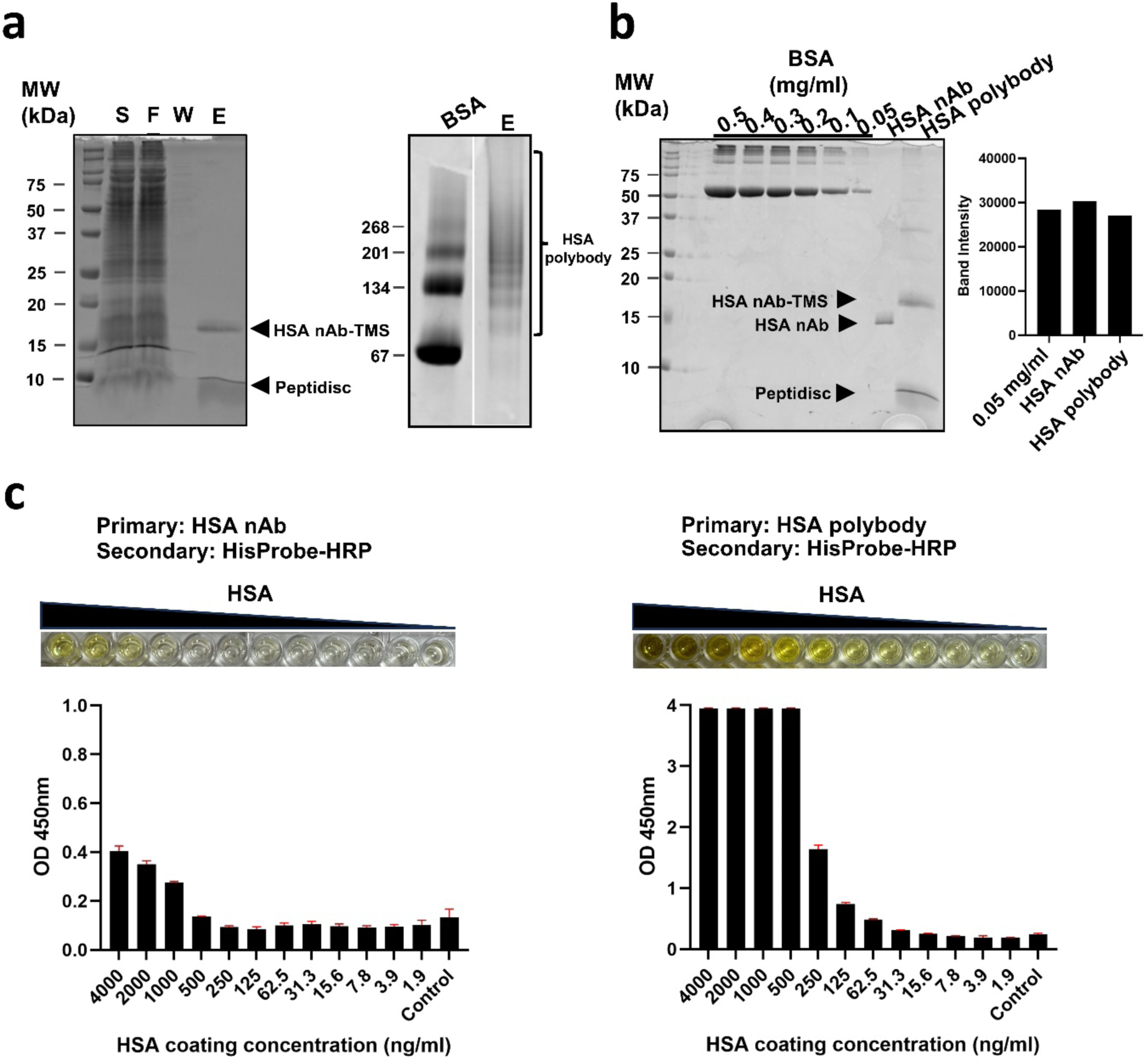
Preparation of the HSA polybody and application to ELISA assays. **(a)** SDS– and native gel analysis of the IMAC-purified and peptidisc reconstituted HSA Pbs. BSA was used as a MW reference on the native gel. S: starting materials (detergent extract); FT: proteins in the flow-through; W: Wash with buffer; E: proteins eluted after peptidisc reconstitution. **(b)** SDS-PAGE analysis of the HSA Nbs and Pbs preparation employed this ELISA assay. The concentration equivalence of the preparations (50 µg/ml) was confirmed by densitometry using BSA as a reference and ImageJ software, then diluted to 1 µg/ml for the ELISA assay application **(C)** Evaluation of the binding affinity of the HSA Nbs and Pbs preparation using indirect ELISA assay. Absorbance was measured at 450 nm. All values are mean ± SD (n=3).

### Development of bispecific polybodies

We next explored whether the composition of the Pbs could be manipulated to achieve a hetero-oligomeric arrangement. To show that Pbs can be made bi-specific, GFP and HSA Nbs were captured in the same Pb. The corresponding Nb-TMS constructs were expressed in *E. coli* and the detergent extracts were mixed at a final 1:1 ratio to favor Pb capture in the same peptidisc, hereafter termed a GFP/HSA Pb. As a control, we mixed the GFP Pbs and HSA Pbs post-reconstitution at a final 1:1 ratio. The SDS-PAGE analysis showed that equivalent amounts of GFP and HSA Nbs were present in these preparations **(Fig. 5a)**. We then assayed the GFP/HSA Pbs bi-specificity using a “sandwich” dot-blot assay **(Fig. 5b)**. In this assay, serial dilutions of the HSA protein (0.26-10 µg/dot) were immobilized onto the nitrocellulose before incubation with the Pbs preparations. After washing, the membrane was incubated with the GFP protein and scanned at 530 nm. As predicted, a fluorescent signal was obtained with the GFP/HSA Pbs, and not with the GFP Pbs or HSA Pbs, either alone or when mixed together. This result demonstrates that GFP and HSA Nbs are present in the same polybody, achieving bi-specificity.

**Figure 5:**
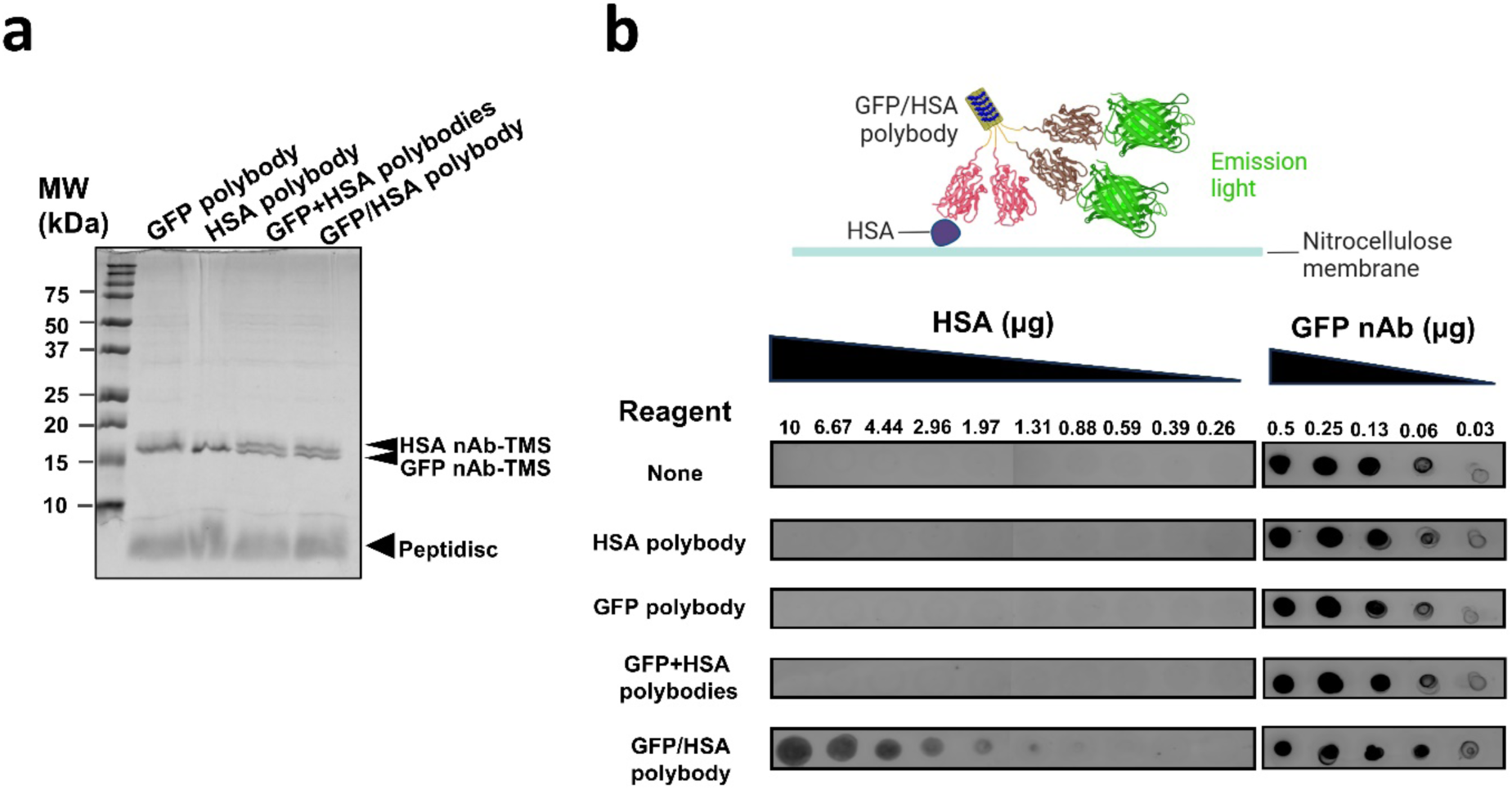
Preparation of a bispecific GFP/HSA polybody and application to dot-blot assays. **(a)** SDS-PAGE analysis of the bispecific GFP/HSA Pbs and comparison to the monospecific version. **(b)** Sandwich dot-blot assay. The quantity of immobilized HSA protein is indicated. As a control, serially diluted GFP Nbs (0.03-0.5 µg/dot) was immobilized on the membrane to ensure consistent exposure conditions across samples. Created with BioRender.com

As an extended application, we created a bi-functional Pbs by co-reconstituting the HSA Nbs with GFP in the same peptidisc, resulting in a fluorescent HSA Pbs. To achieve this, a fusion protein GFP-TMS (31.8 kDa) was constructed (**Supplementary Figure 2**). This engineered GFP-TMS was self-associating into oligomers, hereafter referred to as poly-GFP. Individually reconstituted poly-GFP and HSA Pbs were mixed and used as a control. The SDS-PAGE analysis showed that equivalent amounts of GFP and the HSA Nbs were present in these preparations **(Fig. 6a)**. As expected, when co-reconstituted with the HSA Nbs, the fluorescent HSA Pbs allowed direct detection of the HSA protein at 530 nm **(Fig. 6b)**, but not when poly-GFP or HSA Pbs were employed alone or mixed together. This indicates the successful establishment of a fluorescent HSA Pbs.

**Figure 6:**
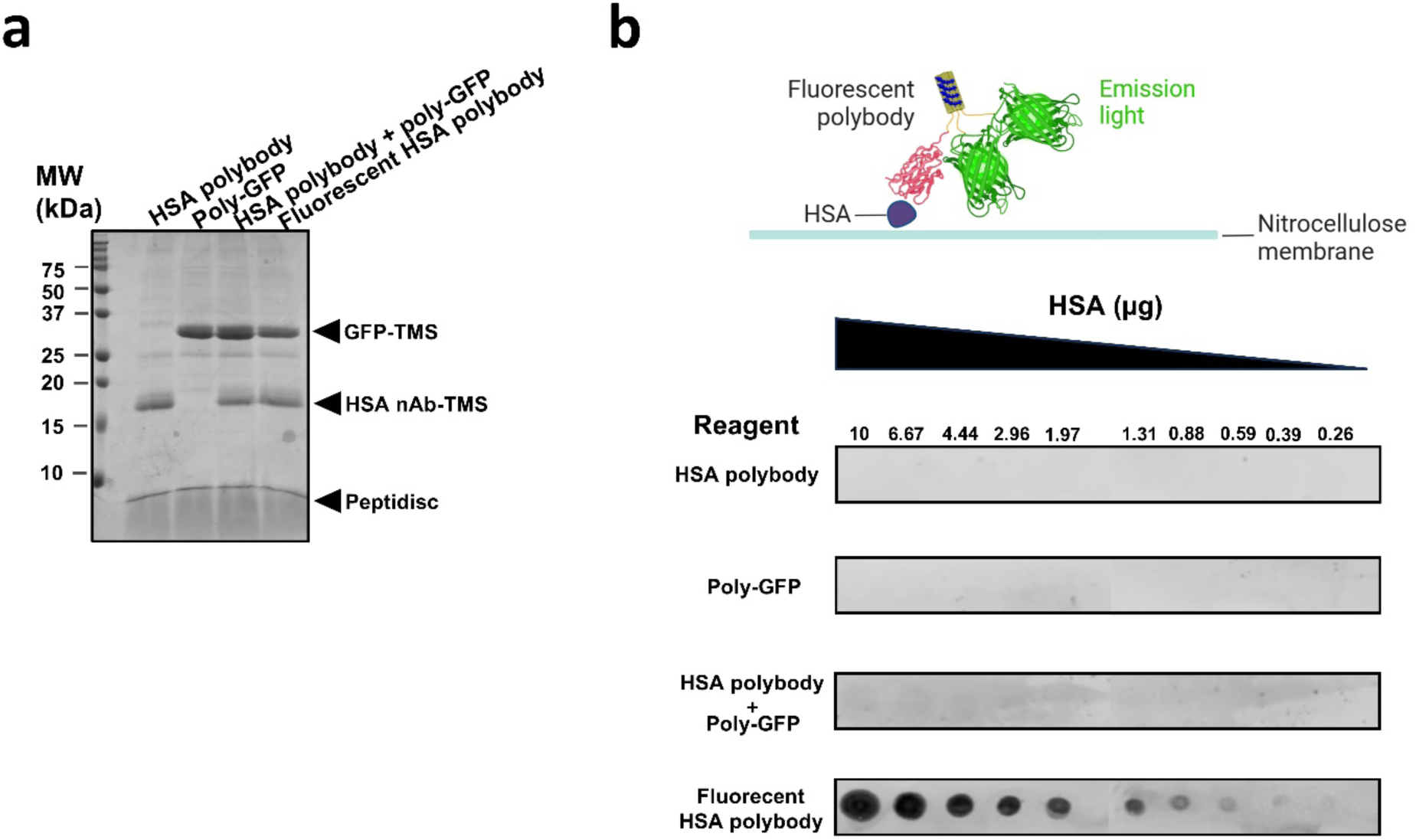
Preparation of a bifunctional polybody comprising the HSA Nb and the GFP protein. **(a)** SDS-PAGE analysis of the fluorescent HSA Pbs and comparison to the standalone version. **(b)** Dot-blot assay. The quantity of immobilized HSA protein is indicated. Created with BioRender.com

## Discussion

We have pioneered an engineering strategy to multimerize proteins based on attaching a hydrophobic transmembrane sequence (TMS) to the C-terminus of a protein of interest. The following hydrophobicity-driven clustering is stabilized with the amphipathic peptidisc scaffold, resulting in the formation of water-soluble protein multimers **(Fig. 1)**. To validate the method from a biological relevant perspective, we used monomeric nanobodies and reformatted them into polybodies. The multimerization was confirmed by native gel, size exclusion chromatography and negative-stain electron microscopy **(Fig. 2**) and the functionality was shown by gel shift assays and biolayer interferometry (BLI). Collectively, the results demonstrated that the polymerization method did not affect the Nb activity since the dissociation constant of monomeric and polymeric GFP Nbs were comparable (2.72 ± 1.60 nM vs 0.75 ± 0.09 nM; **Fig. 3a,b)**. Notably, however, the Pbs dissociation kinetics, determined using an immobilized GFP as the ligand, revealed near-zero dissociation rates **(Fig. 3c)**. Accordingly, multimerization of Nbs results in a cumulative affinity effect, known as avidity, which results in an increased retention of Nbs on their target ^33,35,43–45^. To showcase a biotechnological advantage conferred by the avidity effect, we employed the moderate-affinity HSA Nb (K_d_ ∼ 430 nM) and reformatted it in Pbs **(Fig. 4)**. As expected, the detection limit in the ELISA assay was greatly improved upon polymerisation, decreasing from 1000 ng/mL to 31.25 ng/mL. This ∼32-fold improvement in detection limit is comparable to other Nbs multimerization techniques, which report enhancements ranging from 24-to 175-fold ^28,32,33^.

As a second biochemical application, we applied our polymerization strategy to produce bi-specific Pbs targeting both GFP and HSA. A sandwich dot-blot assay revealed a positive detection signal only when the GFP and HSA Nbs were co-assembled in the same Pb, and not when the individual GFP and HSA Pbs were mixed post-reconstitution **(Fig. 5b)**. The Pbs were stable since their constitutive Nbs did not exchange during the prolonged dot-blot incubations. We further expanded our method by co-reconstituting HSA Nbs with GFP to create fluorescent Pbs. In that case, a fluorescent signal was readily detected upon binding of the Pbs to the HSA protein **(Fig. 6b)**, which may be advantageous when other chemical labeling strategies disrupt the Nb binding site. Finally, based on the same auto-assembly principle, we produced GFP oligomers (termed poly-GFP, **Supplementary Figure 2**), showing that the method can apply to proteins of different size or fold.

Most self-assembly methods, to cite a few – cartilage oligomeric matrix protein, C4bp, p53 tetramerization domain, tetrabrachion, vasodilator-stimulated phosphoprotein, antibody-Fc region, streptavidin, verotoxin 1 B-subunit (VT1B), ferritin, and lumazine synthase ^7,26–29,32–34,45,46^ – are constrained to the production of either multimeric or multispecific Nbs, but not both. Our approach thus expands the current Nb engineering toolbox by offering the option of generating both multimeric and multispecific protein entities using the same strategy. We acknowledge however that our method has limitations, in part due to the uncontrolled polymerization step, producing particles containing variable subunit numbers. This variability, although advantageous in applications where association depends on non-fixed stoichiometry or when the production of oligomers of the desired size is not easily achievable, impedes the precision of the affinity measurements. To address this limitation, fractionating the multimerized protein preparation could help isolate species with the desired copy numbers. Alternatively, changing the primary sequence of the TMS could alter oligomerization kinetics, resulting in a more homogeneous population. Studies have shown that lipids can also influence the oligomeric assembly of membrane proteins by altering oligomerization dynamics ^47,48^. Given the key role of the hydrophobic sequence in our approach, adding exogenous lipids to control Pbs formation represent another interesting optimization step. Finally, we used a C-terminally located TMS to bypass the Sec machinery and mimic the topology of tail-anchored proteins ^49–52^, but other membrane protein topologies could be explored to optimize the production process.

## Statements & Declarations

### Author contributions

Y. C. formal analysis; Y. C. investigation; Y. C. and F. D. v. H. methodology; Y. C. validation; Y. C. data curation; Y. C. and F. D. v. H. conceptualization; Y. C. writing–original draft; Y. C. and F. D. v. H. writing–review and editing; Y. C. software; F. D. v. H. resources; F. D. v. H. funding acquisition; F. D. v. H. project administration; F. D. v. H. supervision.

### Supporting Information

Raw data of bio-layer interferometry analysis (Supplementary Figure 1); measured association and dissociation rates and calculated dissociation constants for the GFP Nb in the avidity-biased measurement (Supplementary Table 1); preparation of a polymeric GFP (Supplementary Figure 2).

## Acknowledgments

We are grateful to Dr. Calvin Yip (University of British Columbia) for providing access to the BLI instrument.

## Data availability

The data that support the findings of this study will be made available by the corresponding author upon reasonable request.

